# Babytwins Study Sweden (BATSS): A multi-method infant twin study of genetic and environmental factors influencing infant brain and behavioral development

**DOI:** 10.1101/2021.04.19.439492

**Authors:** Terje Falck-Ytter, Linnea Hamrefors, Monica Siqueiros Sanchez, Ana Maria Portugal, Mark Taylor, Danyang Li, Charlotte Viktorsson, Irzam Hardiansyah, Lynnea Myers, Lars Westberg, Sven Bölte, Kristiina Tammimies, Angelica Ronald

## Abstract

Twin studies can help us understand the relative contributions of genes and environment to phenotypic trait variation including attentional and brain activation measures. In terms of applying methodologies like electroencephalography (EEG) and eye tracking, which are key methods in developmental neuroscience, infant twin studies are almost non-existent. Here we describe the Babytwins Study Sweden (BATSS), a multi-method longitudinal twin study of 177 MZ and 134 DZ twin pairs (i.e. 622 individual infants) covering the 5 - 36 month time period. The study includes EEG, eye tracking and genetics, together with more traditional measures based on in-person testing, direct observation and questionnaires. The results show that interest in participation in research among twin parents is high, despite the comprehensive protocol. DNA analysis from saliva samples was possible in virtually all participants, allowing for both zygosity confirmation and polygenic score analyses. Combining a longitudinal twin design with advanced technologies in developmental cognitive neuroscience and genomics, BATSS represents a new approach in infancy research, which we hope to have impact across multiple disciplines in the coming years.

## 1. INTRODUCTION

In this article, we describe the multi-modal longitudinal twin study BATSS (**Babytwins Study Sweden)**. The study is based on the idea that combining the field of developmental cognitive neuroscience and twin research can advance knowledge about the origins of individual variation in brain and behavioral development in infancy. While there are existing studies using traditional psychological methods such as questionnaires and direct observation of infant twins, there are no infant (< 2 years) twin studies using a full array of state of the art, advanced methodology from developmental cognitive neuroscience, such as eye tracking and electroencephalography (EEG). These approaches can reveal unique aspects of infant brain and behavioral development.

The study of individual differences continues apace within the field of developmental science. Partly, this is due to the recognized need to predict who is likely to experience certain later outcomes (e.g. disease or developmental difficulties). Key constructs such as equifinality, multifinality, developmental cascades/pathways and probabilistic epigenesis all relate to individual differences in development (Cicchetti & Rogosch, 1996; Gottlieb, 2007; Waddington, 2014). In parallel to the theoretical developments, the gradual accumulation of reliable and valid psychological measures for infancy research has also contributed to the enhanced focus on individual differences in developmental science.

In older children and adults, twin studies have for decades been instrumental for our understanding why people differ in their traits and disease and disorder outcomes (Polderman et al., 2015). By comparing the degree of similarity in MZ and DZ twins, one can quantify the relative contribution of genetic and environmental influences to single phenotypes. Furthermore, the multivariate extension of twin analyses allow for an array of analyses that explore, for example, the degree to which the same genetic influences and the same environmental influences affect two or more phenotypes or explain patterns of stability and change of the same phenotype over development. Twin studies can also shine light on the “heritability of the environment” (i.e. when aspects of one’s environment are shaped by one’s genetic predispositions). Specifically, environments that are separate to each child can be incorporated into the twin design, and thus the role of genetic and environmental influences on a putative environment, such as stressful life events, can be estimated (Shakoor et al., 2016).

In terms of the existing twin literature applying brain-based measures from developmental cognitive neuroscience, EEG studies of infant twins are nearly non-existent. EEG measures electrical activity arising from synchronous neural activity in the brain from sensors placed on the skull. EEG studies have the potential to characterize early neurodevelopmental processes, and unlike eye tracking or other methods, it does not require any overt behavioral response. While EEG has limited spatial resolution, its temporal resolution is excellent, which is used in both time- and frequency-based analyses.

The lack of infant twin EEG studies is surprising given the widespread use of this method in developmental science generally, and the potential value of using non-invasive brain measures to identify heritable psychological traits associated with later neurodevelopmental conditions (e.g. autism; Bosl, Tager-Flusberg, & Nelson, 2018). Studies of older children have found that EEG measures such as frequency band amplitude are highly heritable (Van Baal, De Geus, & Boomsma, 1996; Zietsch et al., 2007). However, to our knowledge, only one twin study of infant EEG exists (Orekhova, Stroganova, Posikera, & Malykh, 2003). This cross-sectional study, which included 95 twin pairs, provided evidence for genetic effects on the alpha frequency band in 7-12 month-olds, but also tentatively suggested that shared environment could play a role for other EEG measures at this age.

Similarly, we are not aware of any eye tracking studies of twins under age of 2 years. Eye tracking uses infra-red light to measure the gaze direction of participants while observing stimuli, Eye tracking has been instrumental to our understanding of young infants’ attentional, perceptual and emerging cognitive abilities, in both the social and non-social domains (Gredebäck, Johnson, & von Hofsten, 2010). Infants develop flexible gaze behavior earlier than many other behaviors (e.g. crawling), and hence, this method provides a unique window into preverbal development. Eye tracking can also be used to assess the size of the pupil, which has proven to be an informative measure as well (Nyström et al., 2018).

One eye tracking study of 2-year-old twins conducted an analysis of eye movement and gaze characteristics in a relatively small sample (83 pairs; Constantino et al., 2017). They found that the tendency to prioritize the eyes versus the mouth during close-up face observation was highly heritable (broad heritability close to 90%). Further, many lower level aspects of eye movements, such as the timing and spatial direction of saccades, were also more similar in MZ than in DZ twins, although caution is needed in light of the small sample size. In light of the extensive use of eye tracking in developmental science generally, and specifically in studies of infants at risk of autism, there is considerable potential to employ this method in studies of young twins.

Among the small number of infant twin studies using methods from cognitive neuroscience, most have used magnetic resonance imaging (MRI; for a review of these twin studies, see Maggioni, Squarcina, Dusi, Diwadkar, & Brambilla, 2020). In terms of differences in brain structure, evidence suggests that genetic factors are important, even very early in life. The magnitude of genetic influence appears to be higher for white matter than grey matter (Gilmore et al., 2010), and region specific (posterior grey matter volume is more heritable than grey matter in frontal areas (Gilmore et al., 2010, see also Duan et al., 2020). Diffusion Tensor Imaging studies suggest significant heritability of early white matter, but substantial regional differentiation, which may reflect the different developmental or maturational timescales of different white matter tracts, for example, in terms of myelination, as well as time periods with more environmental susceptibility (Geng et al., 2012; S. J. Lee et al., 2017; Sadeghi, Gilmore, & Gerig, 2017). To our knowledge, only one study has investigated functional MRI in young twins (Gao et al., 2014).

In contrast, there is a larger body of literature of infant twin studies assessing phenotypes such as temperament, sleep and cognition through questionnaires and observational measurements. Studies focused of language and communication in late infancy and toddlerhood show that shared environment accounts for most of the variance (∼60%) for measures including imitative ability (McEwen et al., 2007), language acquisition (Harlaar, Hayiou-Thomas, Dale, & Plomin, 2008), vocabulary (Hayiou-Thomas, Dale, & Plomin, 2012; Van Hulle, Goldsmith, & Lemery, 2004), and early verbal ability (Galsworthy, Dionne, Dale, & Plomin, 2000). Modest heritability is shown for these type of measures in infancy and early childhood (Dale, Dionne, Eley, & Plomin, 2000; Ganger, 1998; Hayiou-Thomas et al., 2012; McEwen et al., 2007; Robinson, 1999; Van Hulle et al., 2004). For non-verbal cognitive ability, most studies report low-to-moderate heritability in toddlerhood and early childhood (Bishop et al., 2003; Cherny et al., 1994; Matheny Jr, 1980; Petrill et al., 1998; Petrill et al., 1997; Plomin et al., 1993).

Studies of infant attachment report that shared environmental factors explain a large portion of the variation in the late infancy and toddlerhood period (Bakermans-Kranenburg, Van Uzendoorn, Bokhorst, & Schuengel, 2004; Bokhorst et al., 2003; Fearon et al., 2006; Roisman & Fraley, 2008). These results contrast with studies focusing on parent child interaction in other contexts than attachment-eliciting situations, which suggest that genetic factors play a substantial role for variation in infant behavior during parent child interaction, with little contribution from shared environmental factors (Deater-Deckard & O’Connor, 2000; DiLalla & Bishop, 1996).

Behavioral genetic research has also included socio-communicative traits. In the last decade, several studies have focused on early emerging behavioral traits linked to autism spectrum disorder (ASD). Overall, such autistic-like traits are moderately-to-highly heritable in toddlerhood (de Zeeuw, van Beijsterveldt, Hoekstra, Bartels, & Boomsma, 2017; Stilp, Gernsbacher, Schweigert, Arneson, & Goldsmith, 2010). Very high heritability estimates have been found for some specific socio-communicative traits such as attention to eyes and mouth in the second year of life (Constantino et al., 2017), as well as for socio-communicative behaviors rated by parents (Hawks, Marrus, Glowinski, & Constantino, 2019), and reciprocal social behavior (Pohl et al., 2019).

Other areas of early psychopathology and neurodevelopment have been investigated. Hyperactivity was highly heritable in a study with preschool children (2-4 years; Price, Simonoff, Waldman, Asherson, & Plomin, 2001) with no influences of shared environment. Similarly, the heritability of parent rated self-control has been estimated to be substantial with little evidence of shared environmental influences in studies with twins of 2 and 3 years of age (Gagne & Hill Goldsmith, 2011; Gagne & Saudino, 2010, 2016).

Emerging ADHD traits can also include executive functioning dimensions of temperament (i.e., effortful control, and attention control). Temperament in infancy research is a broad but widely studied phenotype. Twin studies suggest that individual differences in infant and child temperament have a moderate to high heritability (Gagne, Vendlinski, & Goldsmith, 2009; Planalp & Goldsmith, 2020; Saudino, 2005), with higher genetic effects for negative aspects of temperament (e.g., anger; Gagne & Hill Goldsmith, 2011) than positive aspects (e.g., positive affectivity and soothability; (Gagne et al., 2009; Goldsmith, Lemery, Buss, & Campos, 1999; Saudino, 2005). Further, motor activity level shows moderate genetic and no shared environment effects (Goldsmith et al., 1999); and significant genetic overlap with ADHD symptoms at 2 years (Ilott, Saudino, Wood, & Asherson, 2010).

Temperament is linked to infant sleep, which is crucial for a healthy development. Sleep has been researched in a few twin studies of infants 6 months and older with moderate sample sizes and parent-reported measures. These studies suggest that early childhood daytime sleep may be driven by shared environmental factors to a substantial extent, whereas the variance in nighttime sleep may be more heritable (Brescianini et al., 2011; Dionne et al., 2011; Fisher, van Jaarsveld, Llewellyn, & Wardle, 2012; Touchette et al., 2013).

Finally, atypical sensory processing (e.g., hyperresponsivity certain sensory signals; Nyström et al., 2018) are increasingly seen as playing a role in a range of neurodevelopmental disorders, including ASD and ADHD (Dunn & Bennett, 2002). However, its genetic and environmental influences early in life have not been extensively investigated, with only one twin study done in 1- to 3-year-olds reporting a moderate heritability for sensory defensiveness and a small contribution from shared environment (Goldsmith, Van Hulle, Arneson, Schreiber, & Gernsbacher, 2006).

Taken together, while several areas of infant and early child development have been the focus of previous behavioral genetic research, there is considerable potential to expand understanding of the genetic and environmental influences on infant development, particularly in the areas of brain, attentional and behavioral development. Furthermore, some of the most influential studies have either begun at 24 months, thus missing the infancy years, or been challenged by small samples of under 300 pairs. With BATSS, we took a multi-method approach to perform deep phenotyping of a wide range of traits at five months and followed a sample of 300 twin pairs longitudinally.

### 1.1 Purpose and aims

The purpose of this article is to describe the BATSS study. We will focus on describing the protocol and procedures, the feasibility of conducting a study of this kind, provide descriptive data about the sample, and discuss putative analysis steps going forward.

## 2. MATERIALS AND METHODS

### 2.1. The classic twin design

BATSS employs a classic twin design. Using twin data, the total variance in a trait can be partitioned into genetic variance, common environmental variance and nonshared environmental variance, which incorporates measurement error. MZ twins share 100% of their segregating DNA and DZ twins share on average 50% of their segregating DNA. All twins who are raised together are assumed to share all of their common environmental influences. Differences identified within MZ twin pairs are explained by nonshared environment. As such, when higher within-pair similarity is observed for MZ than DZ twins, this pattern is assumed to be due to the role of genetic influences on the trait under investigation (Polderman, Benyamin et al., 2015.). In BATSS, the pre-specified target sample size was 620 individuals (310 pairs) based on the size of previous twin studies of toddlers (e.g. Ronald, Edelson, Asherson, & Saudino, 2010). Based on previous twin studies in Sweden, we expected roughly equal numbers of MZ and DZ same sex twin participants

### 2.2. Study protocol

The study protocol is provided in **Table 1**, which also includes full names of measures and abbreviations. Many of the experiments and instruments align with a longitudinal study of infants at elevated likelihood of ASD/ADHD (Jones et al., 2019). The main rationale behind this is to be able to assess the heritability of putative early ASD/ADHD markers (e.g. Nyström, Jones, Darki, Bölte & Falck-Ytter, 2020). In addition, specific measures were added for this study, namely the ITC, SDQ, RBQ, three eye tracking tasks (emotional faces, dynamic singing faces, ANS) and one EEG task (form motion). Many eye tracking and EEG tasks focus on social perception/attention, which is motivated by the abovementioned link to ASD, by the fact that many of these skills/processes emerge early in infancy, and because in older participants, evidence suggest that such socio-cognitive processes may be linked to unique etiological factors, not shared with other aspects of cognition (Shakeshaft & Plomin, 2015).

**Table 1.** BATSS protocol.

|  | Construct/domain | 5 m. | 14 m. | 24 m. | 36 m. |
| --- | --- | --- | --- | --- | --- |
| **Questionnaires** |  |  |  |  |  |
| **Demographics** | Family background | X |  |  |  |
| **Infant-Toddler Sensory Profile** (ITSP; Dunn, 2002) | Sensory processing | X |  |  |  |
| **Medical and Psychiatric History** | Familial risk | X |  |  |  |
| **Communication Development Inventory** (CDI; Fenson et al., 1993; 5-14 "words and gestures", 24-36: "words and sentences") | Language/communication | X | X | X | X |
| **Infant Behavior Questionnaire / Early Childhood Behavior Questionnaire** (short forms) (IBQ-SF; Gartstein & Rothbart, 2003; Rothbart, 1981) / **ECBQ-SF**; Putnam, Gartstein, & Rothbart, 2006). | Temperament | X | X | X | X |
| **Sleep and Settle Questionnaire / Child Sleep Habits Questionnaire-Toddler** (SSQ; Matthey, 2001) / **CSHQ-T**; Owens, Spirito, & McGuinn, 2000; at 24-36 even parts of **Extended Brief Infant Sleep Questionnaire** (EBISQ; Sadeh, Mindell, Luedtke, & Wiegand, 2009) | Sleep | X | X | X | X |
| **Infant Toddler Checklist** (CSBS DP ITC; Wetherby et al., 2004)) | Social communication |  | X |  |  |
| **Quantitative Checklist for Autism in Toddlers** (Q-CHAT; Allison et al., 2008) | Autistic traits |  |  | X | X |
| **Repetitive Behaviors Questionnaire - 2** (RBQ-2; Leekam et al., 2007) | Autistic traits |  |  | X | X |
| **Strengths and Difficulties Questionnaire** (SDQ; Goodman, Ford, Simmons, Gatward, & Meltzer, 2000)) | Psychopathology |  |  | X | X |
| **Vineland Adaptive Behavior Scales** (VABS; Sparrow, Cicchetti, & Balla, 2005)) | Adaptive functioning | X |  |  |  |
| **In-person behavioral assessments** |  |  |  |  |  |
| **Mullen Scales of Early Learning** (MSEL; Mullen, 1995)) | General development | X |  |  |  |
| **Parent-Child Interaction** (videotaped play session; Pijl et al., 2021)) | Social interaction | X |  |  |  |
| **Physical measures:** Head circumference/height/weight | Physical growth | X |  |  |  |
| **EEG/ERP** |  |  |  |  |  |
| **Form/Motion task** (global form and motion coherence perception; Nystrom, Jones, Darki, Bolte, & Falck-Ytter, 2021)) | Visual perception | X |  |  |  |
| social/non-social videos (Jones et al., 2019)) | Functional “resting state” brain activation | X |  |  |  |
| **Eye-tracking** |  |  |  |  |  |
| **Approximate Number System** (ANS; Schröder, Gredebäck, Gunnarsson, & Lindskog, 2020)) | Cognition | X |  |  |  |
| **Dynamic faces** | Attention to social stimuli | X |  |  |  |
| **Emotional Faces** | Attention to social stimuli | X |  |  |  |
| **Face pop out** (Elsabbagh et al., 2013) | Attention to social stimuli | X |  |  |  |
| **Gap attention-shifting task** (Johnson, Posner, & Rothbart, 1991) | Disengagement of attention | X |  |  |  |
| **Naturalistic Scenes** (de Urabain, Nuthmann, Johnson, & Smith, 2017; Jones et al., 2019) | General attention | X |  |  |  |
| **Pupillary Light Reflex** (Nystrom, Gredeback, Bolte, & Falck-Ytter, 2015; Nyström et al., 2018) | Visual sensory processing | X |  |  |  |
| **DNA and other biological samples** |  |  |  |  |  |
| **Hair and Nail Samples** | Biological processes | X |  |  |  |
| **Saliva** | DNA | X |  |  |  |
| **Questionnaires about the parents’ behavior** |  |  |  |  |  |
| **Autism quotient** (AQ; Baron-Cohen, Wheelwright, Skinner, Martin, & Clubley, 2001) | Autistic traits |  | X |  |  |
| **ADHD self report scale** (Scale_(ASRS-v1.1), retrieved from https://nyulangone.org/files/psych_adhd_checklist_0.pdf) | ADHD traits |  | X |  |  |

As many of the tasks and measures are well established, we do not describe them all in detail here, but additional information can be found elsewhere Jones et al. (2019) or in the original works cited in **Table 1**. For other measures not described in detail elsewhere, we provide a brief description here.

#### Demographics and Medical/Psychiatric History

At the 5 month visit, we administered questionnaires and conducted interviews about basic background information such as date and place of birth, languages spoken in home, occupational and educational status and family income. Information about the twins’ medical conditions was also collected (including twin specific birth complications), as well as the presence of developmental and psychiatric conditions in family and relatives. A set of items surveyed parents’ perceived similarity of their infant twins.

#### Eye tracking tasks

(not described elsewhere). *Dynamic faces*. This eye tracking task consisted of dynamic faces (n = 20, duration 4-12 seconds), divided into three conditions: singing, non-singing nursery rhymes, silent. This task measures infants’ preference for eyes versus mouth (Constantino et al., 2017; Hunnius & Geuze, 2004) in the context of naturalistic face stimuli. This task was designed specifically for BATSS, but stimuli properties (e.g. duration) was similar to what has been used previously with infants. *Emotional Faces*. This task showed triplets of static faces in each stimulus (n = 12, duration 5 seconds), of which two were neutral and one had an emotional expression (happy). This task is intended to capture individual differences in attention to emotional facial stimuli, and was adapted from a longitudinal study of social attention.

#### Biological samples

*Saliva*. Saliva samples were collected by research assistants from all the infant twins using the DNA Genotek OG-575 (DNA Genotek Inc.) collection kit during the study visit. *Hair*. A lock of hair was cut from the posterior vertex region of the head by parents, as close to the scalp as possible. *Nails*. Sample of at least 4mm in length were collected from fingernails and toenails. Hair and nail samples were collected to assess metabolic changes over time related to prenatal and early postnatal exposures to metals and/or other substances.

### 2.3. Recruitment and Testing Procedure

#### 2.3.1. Recruitment and first telephone interview

Same-sex twin families were identified via the Swedish Population Registry (Folkbokföringsregistret, hosted by the Swedish Tax Agency). In total, 1068 families with same-sex twins were invited to join the study via letters. If families agreed to participate during a follow-up telephone call, a screening interview was performed. To be included in the study, the twins had to hear Swedish at home from at least one parent and live with at least one biological parent. Parents also needed to be willing to share information about medical and psychiatric history in the family, demographic background and delivery. Reasons for exclusion were hearing or vision impairments, premature birth (defined as prior to week 34 of gestation), epilepsy or seizures, medical conditions that were likely to affect brain development, ability to participate in the study, and the presence of known genetic syndromes. In a few cases, we included and tested twins with ambiguous medical conditions (12 pairs with twin to twin transfusion syndrome reported by parents, one twin with low birth weight, one twin with spina bifida and two twins with seizures at birth; these are included in the total sample described in this article).

#### 2.3.2. Five month in-person visit at the lab

At age 4-5 months, a letter was sent home to the parents with information about the visit, printed questionnaires and directions for completing online questionnaires (see **Table 1** for overview).

The 5 month visit was typically performed in one day, lasting approximately five hours. At least two research assistants were present, each having the responsibility for specific tasks. During the visit, the twins were called by their actual names in communication with their parents and each twin had their own personal ‘schedule’. This enabled the research assistants to keep track of the tasks performed and minimized the risk of mixing the twins up. If the research assistants had difficulty telling them apart, due to the twins being very alike and/or dressed similarly, stickers with the numbers ‘1’ and ‘2’ were placed on their abdomens. The twins performed different tasks at the same time and we generally encouraged both parents to be present during the visit, so that each child could be accompanied by one caregiver in all tasks. When it occasionally was not possible for both parents to attend the visit (i.e., only one parent being able to take leave from work or in single parent households), we had a third research assistant present acting as a ‘stand in’. If this was the case, we always informed the parent(s) prior to the visit, to verify that this was acceptable to them. Occasionally, other relatives and friends could be present and to assist in such cases.

We started with performing the tasks that were most demanding for the infants, namely EEG (approx. 11 minutes), and the Mullen Scales of Early Learning (MSEL; approx. 10-15 minutes). Later in the visit, we performed Eye Tracking (several experiments in two separate sessions; approx. 6 and 10 minutes respectively) and Parent-Child Interaction (PCI), a playtime with the mother and one twin at a time that was video recorded for subsequent behavioral coding (approx. 15 minutes). However, the order of tasks was not entirely fixed and varied depending on the moods and needs of the twins.

We took breaks when needed, to let the twins rest and eat. In the middle of the day, we had a longer break for lunch and carried out parent report interviews (about demographic details such as education and employment, as well as medical and psychiatric history in relatives, see Table 1). The parents received gift vouchers as compensation for participating.

#### 2.3.3. Follow up questionnaire packages (14-36 months)

The twins were followed up through online questionnaires at 14, 24 and 36 months (see **Table 1** for full overview of included questionnaires). At these ages, we sent a letter with detailed instructions about the questionnaires and login information, together with a gift voucher. If the questionnaires were not completed after two weeks, we sent a brief reminder via email. If the questionnaires were not completed after a month, we initially sent a reminder letter, but this routine was later changed to a phone call to assist the parents with any questions or technical issues. The last date for the parents to complete the questionnaire before the time point was considered passed was at 15 months for the 14 months time point and 30 months for the 24 months time point; for the 36 months time point there was no last date for completion. The exact time point of questionnaire competition was recorded, allowing for further age-based selection/analysis if needed.

#### 2.3.4. Sample collection and Genotyping

Saliva samples were collected from all the infant twins using the DNA Genotek OG-575 collection kit during the study visit. All saliva samples were stored and processed for DNA extraction at the Karolinska Institutet Biobank. DNA extraction was done using Chemagen kits based on magnetic bead separation in using the automated Hamilton ChemagicSTAR® platform. DNA quantity and quality was checked before proceeding to genotyping. The twins were genotyped using the Illumina Infinium Global Screening Array version 3 Infinium Assay using standard protocols at the SNP&SEQ Technology Platform at Uppsala University.

## 3. RESULTS

### 3.1. Recruitment summary

In total, 1068 families were invited to participate in BATSS. Because we recruited from the Swedish Population Registry, these individuals represent most same-sex twins born in the targeted area during the period in question (**Supplementary Figure 1)**. In total, 46.9% of the invited families responded to our recruitment materials and could be interviewed by telephone. A total of 567 (53.1%) of the invited families did not participate, even though brief reminder letters were sent to most of them. **Figure 1** outlines the categories of reasons given by families who did not participate.

**Figure 1.**
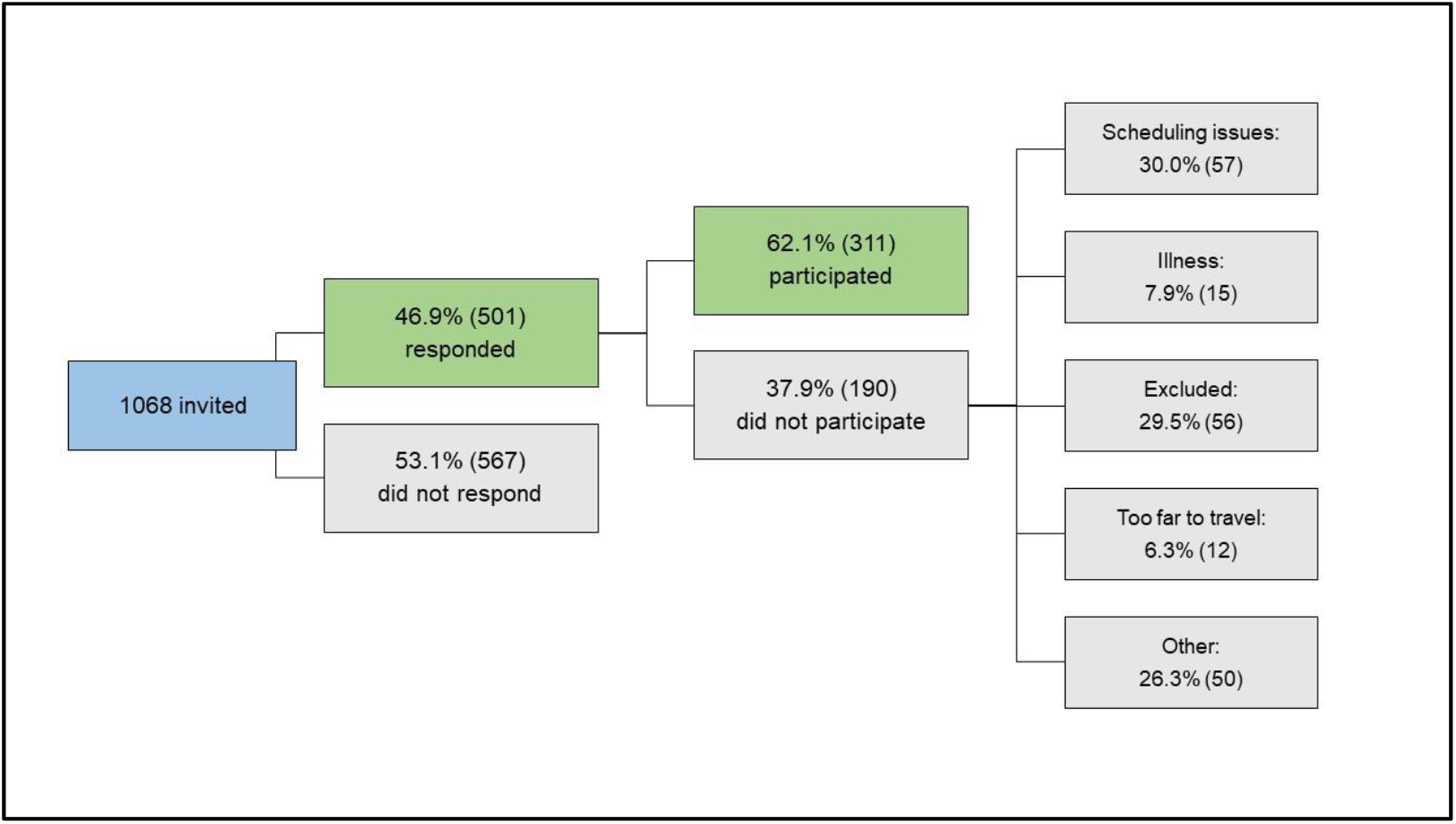
Overall recruitment statistics.

### 3.2. Sample Characterization

**Table 2** summarizes the main characteristics of the final BATSS sample, as well as MZ and DZ subsamples. As can be seen, the total sample consists of 311 twin pairs, of whom 177 are MZ and 134 are DZ. All twin pairs in the BATSS sample were same-sex, with 151 pairs being female and 160 male. All pairs lived in the greater Stockholm and Uppsala area in Sweden upon recruitment, with 27.6% living in central Stockholm. **Supplementary Table 1** summarizes response rates for online questionnaires.

**Table 2.** Participant characterisation^1^.

|  | Total Sample | Subsample: MZ | Subsample: DZ |
| --- | --- | --- | --- |
| **N (pairs)** | 311 | 177 (56.91%) | 134 (43.09%) |
| **N female** | 151 (48.55%) | 84 (47.46%) | 67 (50.00%) |
| **Age (days)<sup>2</sup>** | 167.54/8.65 [145-203] | 167.47/8.49 [149-194] | 167.63/8.87 [145-203] |
| **Parental education<sup>3</sup>** | 3.29/0.85 [1-4] | 3.27/0.86 [1-4] | 3.33/0.84 [1-4] |

<sup>1</sup> n for individual variables vary slightly (298-311 pairs).
<sup>2</sup> Age at 5 months assessment. Mean/sd [min max]; this format applies generally unless otherwise specified.
<sup>3</sup> Education level on a scale from 1-4, where 1 = Primary, 2 = Secondary, 3 = Undergraduate ( $\leq 3$ years) and 4 = Postgraduate level ( $> 3$ years).

|  |  |  |  |
| --- | --- | --- | --- |
| **Family income<sup>4</sup>** | 6.57/2.32 [1-10] | 6.49/2.24 [1-10] | 6.70/2.42 [1-10] |
| **Past or current psychiatric condition in first degree relative<sup>5</sup>** | 31.83% | 32.49% | 30.97% |
| **Both parents born in Sweden (At least one parent born in Sweden)** | 73.14% (89.71%) | 75.00% (91.53%) | 70.68% (87.31%) |
| **Gestational age<sup>6</sup>** | 36.70/1.20 [33-39] | 36.51/1.20 [33-39] | 36.94/1.15 [34-39] |
| **Birth weight (grams)** | 2643/426 [1350-4000] | 2574/404 [1520-3525] | 2733/437 [1350-4000] |
| **C-section** | 53.86% | 51.90% | 56.52% |
| **Maternal smoking 1st trimester and the month before (2nd-3rd trimester)** | 3.62% (0.33%) | 4.57% (0.57%) | 2.33% (0.00%) |

<sup>4</sup> Family income per month. Scale 1-10 where 1 = <20K, 2 = 20-30K, 3 = 30-40K, 4 = 40-50K, 5 = 50-60K, 6 = 60-70K, 7 = 70-80K, 8 = 80-90K, 9 = 90-100K and 10 = >100K (SEK).
<sup>5</sup> Reported past/current ASD, ADHD, intellectual disability, panic or anxiety disorder, depression, bipolar disorder, schizophrenia and/or admittance to psychiatric care of mother, father or full siblings of the whole sample. Classified binarily (1 = yes to at least one, 0 = no to all).
<sup>6</sup> Gestational age of the whole sample as defined by the number of completed gestational weeks. One pair was born in week 33+6 (weeks+days).

To further assess the representativeness of our sample, we compared socioeconomic status of our sample with the population of Stockholm and Sweden as a whole, using statistics from Statistiska Centralbyrån (‘Statistics Sweden’, SCB; **Table 3**).

**Table 3.** Socioeconomic status.

|  | Education<sup>1</sup> |  |  | Income<sup>2</sup> |  |  |
| --- | --- | --- | --- | --- | --- | --- |
|  | Primary | Secondary | Tertiary | Low | Medium | High |
| **BATSS** | 0.96% | 15.06% | 83.97% | 19.34% | 54.01% | 26.64% |
| **Stockholm<sup>3</sup>** | 6.97% | 28.30% | 64.73% | 44.19% | 38.74% | 17.08% |
| **Sweden<sup>3</sup>** | 8.53% | 34.17% | 57.30% | 51.31% | 38.97% | 9.72% |

<sup>1</sup>Mothers' highest level of completed education (primary ( $\leq 10$ years of education on ISCED levels 1-2), secondary ( $\leq 3$ years of education on ISCED levels 3-4) and tertiary ( $> 0$ years of education on ISCED levels 5-8)). Education levels represent the ISCED levels of education (UNESCO Institute for Statistics, 2012).
<sup>2</sup>Mothers' yearly income (low: $\leq 299$ K SEK, medium: 300-499K SEK and high: $\geq 500$ K SEK), mean age 34 years.
<sup>3</sup>For comparison, we have included data showing the income and education of the population of Stockholm county and Sweden, compiled 2019 (women aged between 25-44 years; source Statistiska Centralbyrån, SCB).

### 3.3. Molecular genetic analyses

#### 3.3.1. Quality control and Imputation

In 117 pairs of MZ twins (whose zygosity was confirmed earlier via a simplified genetic analysis; (Hannelius et al., 2007)), only one of the infants in the pairs were genotyped with the Illumina Infinium array. This array consists of 730 059 markers of which 97.8% had sample call rate 98% with average SNP call rate per sample 99.3%. Quality control (QC) for the raw genotyping at the individual and marker level was completed using PLINK v1.90. Thereafter, we imputed both autosomal and X chromosomes using IMPUTE2 followed with additional QC There were 518 570 markers remaining after genotyping QC and 7 507 876 markers after imputation QC.

#### 3.4.2. Polygenic scores

Polygenic scores were calculated using PRS-CS (Ge, Chen, Ni, Feng, & Smoller, 2019) for a range of development-related and psychopathological traits (ADHD, Demontis et al., 2019; ASD, Grove et al., 2019; Bipolar disorder, Stahl et al., 2019; Major depressive disorder, Howard et al., 2019; Schizophrenia, Ripke, Walters, O’Donovan, & Consortium, 2020; IQ, Savage et al., 2018); Educational attainment, Lee et al., 2018; Physical Height, Yengo et al., 2018). The distribution of the different polygenic scores and the correlational pattern between them were expected and are shown in **Supplementary Figure 2**. Genetic ancestry can affect the validity of polygenic score analyses, because the base rate of specific SNPs can vary across populations. To control for this one needs to quantify genetic ancestry. Therefore, a principal component analysis (PCA by EIGENSOFT 7.2.1) was performed based on the BATSS and two reference datasets (HapMap Phase III (HapMap3), representing individuals with genetic ancestry from Asia, Africa and Europe and, SWEGEN, representing the genetic ancestry of the Swedish population (Ameur et al., 2017; Gibbs et al., 2003) to analyze population stratification of our twin sample in comparison to the reference datasets (using *R* 3.6.3). Visual inspection of the results of the PCA confirmed that the sample was largely homogenous in terms of Swedish/European ancestry, with only a minority of mixed genetic ancestry or non-European genetic ancestry (for comparison, see questionnaire-based data in **Table 1**)

## 4. DISCUSSION

The main aim of this article was to describe the BATSS study, focusing on its methodology, feasibility and participant characteristics. While, as reviewed in the Introduction, previous research studies have included infant twins, very few studies have taken a multi-method approach and there are no well-powered twin studies employing eye tracking and EEG. Further, few twin studies of infants have included a comprehensive, longitudinal protocol spanning behavior and brain measures and covering psychological constructs from early basic perception to later language skills. By introducing eye tracking and EEG methodologies, the current study can help us understand the etiological factors behind a range of informative processes ranging from low level perception and attention to complex behaviors. The multi-level, longitudinal protocol, which includes both these novel methods and more classical assessment approaches, has the potential to provide a more complete picture of cognitive and brain development than has been possible before.

The study shows that it is possible to recruit a relatively large number of infant twin families and assess them at a steady pace within a medium-sized metropolitan area (greater Stockholm area ∼2 million people; **Supplementary Figure 1**). The relatively high final inclusion rate (approximately 29% of the population in the targeted area; **Figure 1**), despite requiring in-person visits to the lab, suggests high motivation among infant twin parents to participate in research in Sweden. It is notable that initially, nearly 50% of the target population were interested in participating (**Figure 1**), but some were ultimately not able due to scheduling issues or not fulfilling inclusion criteria. In addition, while 53.1% of the targeted population did not respond to our materials, this does not necessarily mean that all these families were negative to participation. Taken together, we conclude that the motivation to participate in research is very high among infant twin parents in Sweden: At least about half of the population is interested in participation in principle. This may reflect general interest in research and an appreciation of the fact that twins are particularly valuable for science (Sweden has a strong tradition of twin research of older children and adults; e.g., (Anckarsäter et al., 2011; Zagai, Lichtenstein, Pedersen, & Magnusson, 2019), but it can also reflect other factors such as generous parental benefits in Sweden, which allow parents to participate with few practical or economical negative consequences.

Importantly, the study shows that it is possible to conduct multi-method assessments within cognitive neuroscience (brain-based measurements, direct behavioral assessments, questionnaires, biosamples) from two infant twins within one day. Again, this may reflect the generous parental benefits in Sweden allowing both parents to be on leave at the same time, and thus both present during testing. However, it is doable in principle to conduct the research with only one parent, and extra research assistants. Taken together, the BATSS study demonstrates high feasibility of comprehensive infant twin studies in other areas with similar societal and geographical characteristics.

There was a slight bias towards MZ twins in the sample (**Table 1**). This may reflect true differences in rates of MZ vs DZ among same-sex twins, but is likely to reflect higher interest in participating in research in MZ twin families. There was a near-equal distribution of females vs males, both in the total sample and in the separate MZ and DZ groups. While the age range (4.8 - 6.7 months, mean 5.5 months) is relatively narrow, we will take age into account in data analyses (e.g. use as covariate). Indeed, this age range is marked by considerable development across many domains of cognitive and brain development.

The MZ and DZ groups were similar in terms of age, and parental education, income, psychiatric family history, and region/country of birth of parents. Because twin analyses build on comparisons between MZ and DZ twins, it simplifies analysis and interpretation that these groups do not differ on background variables. Further, MZ and DZ groups were similar in terms of gestational age and frequency of C-section. MZ twins were around 200g lighter at birth compared to DZ (P=.001). While this difference was marginal at the descriptive level, it could be important to include birth weight as a covariate in analyses.

Compared to the mean of the general Stockholm population, the sample had a higher average level of formal education and somewhat higher average income (**Table 3**). This difference is consistent with studies of older twins in Sweden (e.g. Taylor et al 2020) and should be taken into account in any findings from the study. Little is known about how SES may moderate heritability estimates in early infancy (for an example of this phenomenon in young children, see (Turkheimer, Haley, Waldron, d’Onofrio, & Gottesman, 2003)) but as with all twin samples, the results should be considered in light of the sample characteristics

Within the framework of the classical twin design, both univariate and multivariate analyses will be conducted. Given the young age of the participants, it is of particular interest to compare heritability and estimates of environmental influence for the different types of measures, to understand which aspects of development are under relatively higher versus lower genetic and environmental influence. Further, extending the classic twin design to include measures of the environment (such as the parent’s behavior during parent-child interaction) will also help us understand to what extent the child’s genetic predispositions affect their social environment (e.g. (DiLalla & Bishop, 1996; Kennedy et al., 2017)). It is also of primary interest to understand the genetic landscape across different phenotypes (multivariate twin modelling) to understand to what extent development within and across domains is influenced by a set of shared versus unique genetic and environmental factors. Ultimately, it will also be beneficial, to both the behavioral genetic and development science fields, to account for our longitudinal design in multivariate models to study genetic and environmental continuity and change in cognitive and brain development across the first few years of life.

In addition to comparing similarity in MZ versus DZ twins (classic twin analyses), we will be able to capitalize on MZ differences within the sample, based on the logic that if MZ twins are different, it must reflect differences in non-shared environment (but see (Jonsson et al., 2021)). This design can be powerful in identifying (or falsifying) potential causal pathways linked to environmental exposures. For example, research has shown atypical concentration of certain metals in the teeth of children with autism already in infancy (Arora et al., 2017). We can use similar analyses to check whether any differences in metal concentration in nails or hair in MZ twins correlate with differences in development within MZ pairs.

The sample has been genotyped and polygenic scores have been calculated for all individuals who provided DNA samples and who passed our quality control stages. Therefore, our study is in a position to be able to contribute to understanding of how the common genetic architecture in neurodevelopmental conditions, such as autism and ADHD, and traits and conditions that affect people in later life, such as educational attainment and schizophrenia, influence cognitive and brain development in the first months of life, long before these conditions or traits have emerged (for a recent example of this approach in infancy research, see Gui et al 2020). Polygenic score analyses will be conducted in the sample and we hope the sample can also be used more broadly to advance understanding of the genetic basis of neurodevelopmental, cognitive and psychiatric phenotypes. At present, molecular genetic research on infancy lags far behind research on older samples (Papageorgiou & Ronald, 2013, 2017); the potential to work with the genetic data within the sample represents another opportunity afforded by the BATSS dataset.

## Acknowledgements

The authors thank all participating families, as well as research assistants Joy Hättestrand, Johanna Kronqvist, Sofia Jönsson, Anna Kernell, Carolin Schreiner, Sophie Lingö, Angelinn Liljebäck, Isabelle Enedahl, Matthis Andreasson, Lisa Belfrage, Mattias Savallampi, Isabelle Ocklind and Hjalmar Nobel Norrman. The authors thank prof Paul Lichtenstein for general support. The work leading to these results were supported by Stiftelsen Riksbankens Jubileumsfond (NHS14-1802:1; Pro Futura Scientia [in collaboration with SCAS]); the Swedish Research Council (2018-06232); Knut and Alice Wallenberg foundation; and the European Union (Initial Training Network BRAINVIEW; 642996). LT and DL were supported by the Swedish Foundation for Strategic Research FFL18-0104 and China Scholarship Council. We acknowledge the KI Biobank for handling the biological samples, SNP&SEQ Technology platform at Uppsala University for genotyping and the Swedish National Infrastructure for Computing (SNIC) at UPPMAX, partially funded by the Swedish Research Council through grant agreement no. 2018-05973” for computations.

## Author contribution

Conceptualization: TFY with contributions from AR, SB, KT; Methodology: TFY, KT, AR, MT, SB, LW; Formal analysis: LH, KT, DL, CW, IH; Data curation: TFY, LH; Investigation: MS, LH, LM; Writing (original draft): TFY with contributions from AR, MS, AMP, LH, CW; Writing (review and editing): all authors; Supervision: TFY with contributions from AR, MT. Visualization: DL, LH, AMP; Project administration: TFY with contributions from LH, MS; Funding acquisition. TFY, AR, KT, SB.

## Supplementary materials

### Supplementary Results

In order to test the polygenic scores predictive power in relation to one reliable and accessible phenotype in our sample, we ran a GEE model to predict physical height of the infant at the time of testing. Although the polygenic score for height is based on adult height, an infant’s height is associated with later height and can therefore be used to indirectly validate these scores (Cole, T. J., & Wright, C. M. (2011); Eide, M. G., Øyen, N., Skjœrven, R., Nilsen, S. T., Bjerkedal, T., & Tell, G. S. (2005). Infant’s height was regressed on sex and age before being included in a model using the PRS for height (regressed on 10 components of ancestry) as predictor, with family ID as a cluster variable. Both variables were scaled before analysis so that the Beta estimate can be interpreted as a correlation coefficient (i.e. association could range from -1 to 1). PRS for height was a significant predictor of height at the time of testing (total n = 594, complete n = 578 /292 clusters; Beta = 0.27, z statistic = 5.184, p < .001).

## Supplementary Tables and Figures

**Supplementary Table 1.** Response rates to the questionnaires packages

|  | 5m questionnaires | 14m questionnaires | 24m questionnaires | 36m questionnaires |
| --- | --- | --- | --- | --- |
| **All completed** | 93% | 75% | 64% | 65% |
| **Partially completed** | 6% | 12% | 12% | 9% |
| **None completed** | 1% | 13% | 24% | 27% |

The 24 and 36 months time points will be completed in 2022 and 2023 respectively, response rates in the table are at the time of writing (March 2021).

**Supplementary Figure 1.**
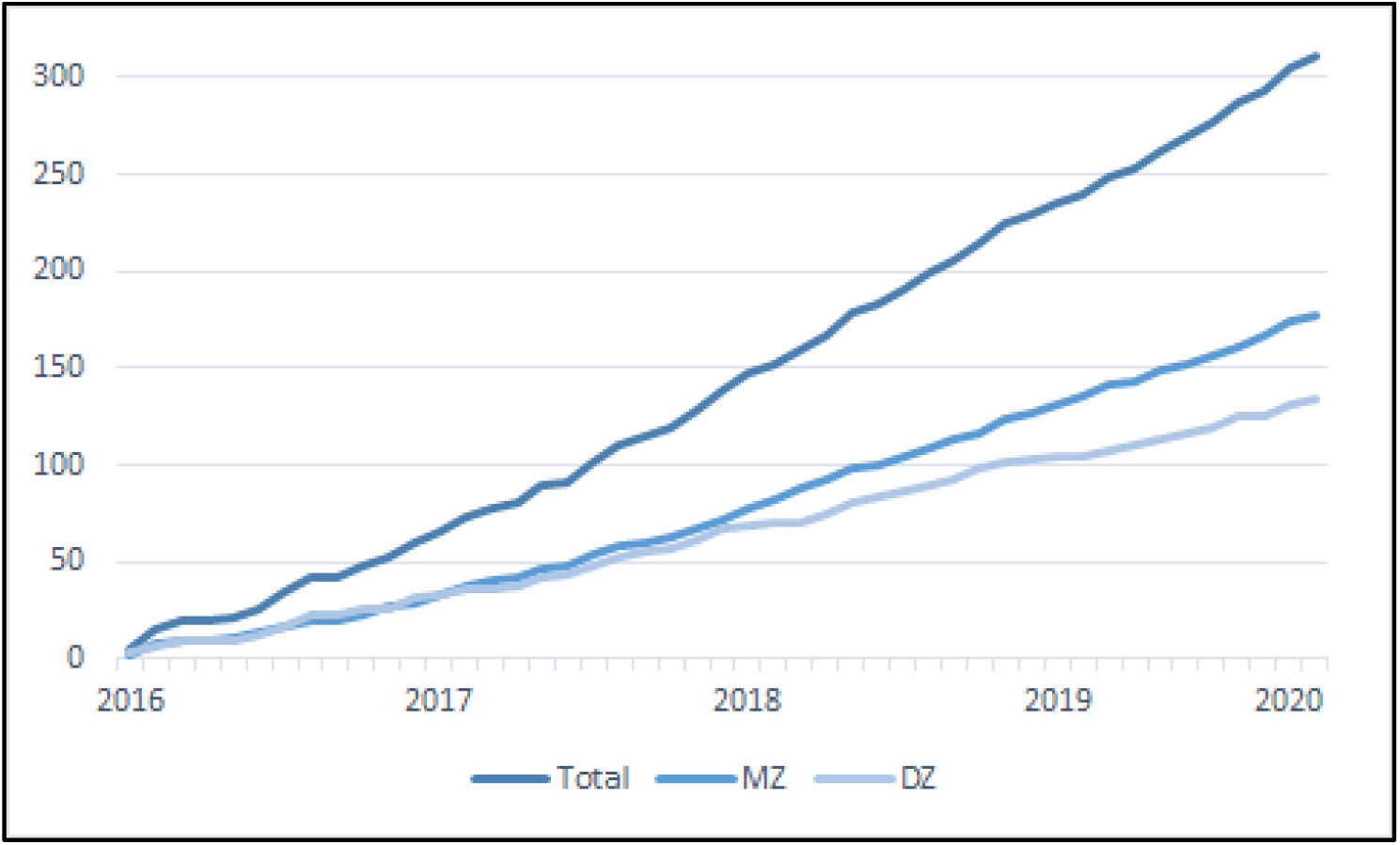
The cumulative number of twin pairs tested per month over the whole study period.

**Supplementary Figure 2.**
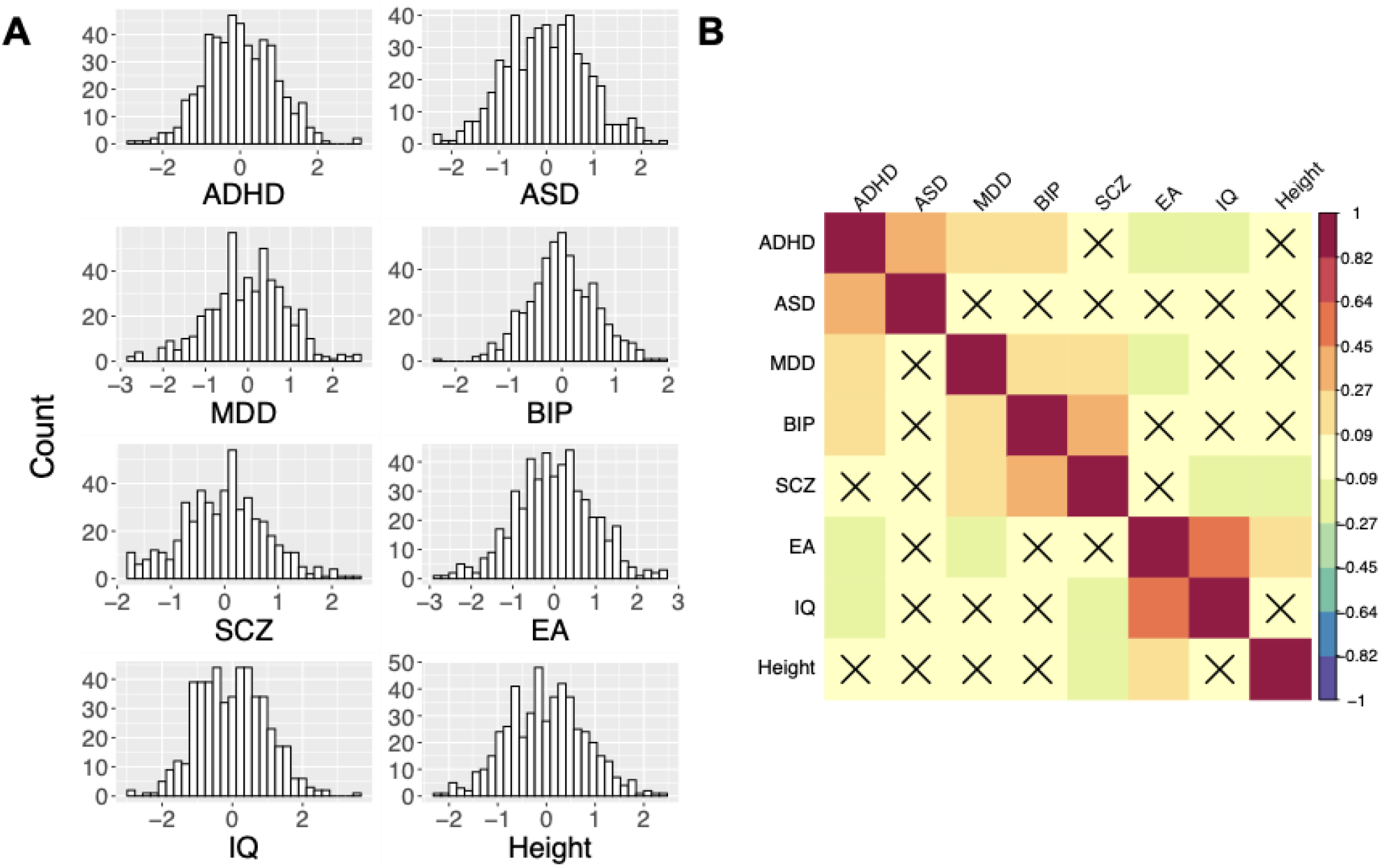
**A**. The distribution of the different polygenic scores **B**. The correlations between the polygenic scores, in the BATSS sample (including uniquely genotyped individuals only, see section Quality Control and Imputation). ADHD = Attention-Deficit Hyperactivity Disorder, ASD = Autism Spectrum Disorder, MDD = Major Depressive Disorder, BIP = Bipolar Disorder, SCZ = Schizophrenia, EA = Educational Attainment, IQ = Intelligence. Scores were regressed on 10 Principal components and scaled before analysis. Color represents strength of the pearson correlation coefficient, cross represents non-significant coefficients (P-threshold is .05).

